# Time cell encoding in deep reinforcement learning agents depends on mnemonic demands

**DOI:** 10.1101/2021.07.15.452557

**Authors:** Dongyan Lin, Blake A. Richards

## Abstract

The representation of “what happened when” is central to encoding episodic and working memories. Recently discovered hippocampal time cells are theorized to provide the neural substrate for such representations by forming distinct sequences that both encode time elapsed and sensory content. However, little work has directly addressed to what extent cognitive demands and temporal structure of experimental tasks affect the emergence and informativeness of these temporal representations. Here, we trained deep reinforcement learning (DRL) agents on a simulated trial-unique nonmatch-to-location (TUNL) task, and analyzed the activities of artificial recurrent units using neuroscience-based methods. We show that, after training, representations resembling both time cells and ramping cells (whose activity increases or decreases monotonically over time) simultaneously emerged in the same population of recurrent units. Furthermore, with simulated variations of the TUNL task that controlled for (1) memory demands during the delay period and (2) the temporal structure of the episodes, we show that memory demands are necessary for the time cells to encode information about the sensory stimuli, while the temporal structure of the task only affected the encoding of “what” and “when” by time cells minimally. Our findings help to reconcile current discrepancies regarding the involvement of time cells in memory-encoding by providing a normative framework. Our modelling results also provide concrete experimental predictions for future studies.

## Introduction

The cognitive ability for biological organisms to remember an episodic event relies on the encoding of “what happened when” by the brain ^1^. Since the discovery of “place cells” ^2^, which are cells that consistently fire when the animal occupies a certain location in space, the hippocampus has long been theorized to contribute to memory by organizing spatial knowledge into a cognitive map. As the temporal analogue of place cells, several recent studies have identified neurons in hippocampus CA1 ^3–8^ and CA3 ^8–10^ that tile the interval between discontiguous events by firing sequentially at successive moments in time, suggesting that these “time cells” support the temporal organization of episodic memory by encoding elapsed time. The subsequent observations of such time cells throughout the brain in multiple mammalian species ^10–18^ confirmed that such a dynamical regime was wide-spread and complementary to the previously reported ramping-based model for tracking time ^19–24^, in which neurons can estimate elapsed time using monotonically increasing or decreasing neuronal firing rates. Interestingly, multiple studies have demonstrated that the same population of hippocampal time cells form distinct sequences during the mnemonic delay following the presentation of different sensory stimuli ^4,5,25^, suggesting a potential mechanism by which the hippocampus integrates information about “what” and “when” as part of the process of encoding episodic memories.

However, there exists some discrepant evidence that makes it unclear whether these time cells are involved in storing memories, or if they are an emergent phenomenon related to non-mnemonic processes. For example, Salz et al.^9^ showed that these time cell sequences emerged from not only the mnemonic delayed alternative task but also a “looping task” that contained no memory load. Sabariego et al.^8^ reported that sequences formed by the hippocampal time cells during a spatial working memory task did not distinguish between different trial types. More recently, Ahmed et al.^26^ showed that CA1 activity patterns were neither consistent nor informative during the delay period in trace fear conditioning. These findings hint at the possibility that the observations of so-called time cells may merely be an epiphenomenon as a consequence of assumption-based analyses, rather than a mechanism by which the brain bridges mnemonic gaps between events. We hypothesize that the discrepancies in current data are a result of different studies using tasks that involve different cognitive demands and different temporal structures, making direct, well-controlled comparisons impossible.

One way to address this is to use computational models of simulated agents, wherein we can both fully control the demands of the task and perform rigorous decoding analyses to investigate the effects of these task-related factors. Thankfully, recent advances in artificial intelligence have shed light on a new path to understanding neuroscience problems ^27,28^. Specifically, artificial neural networks (ANNs) trained with appropriate loss functions and architectures provide a means to explore normative and functional hypotheses about neural computation at the algorithmic-level. Reinforcement learning (RL), in particular, is well-suited to modelling real neural computation thanks to the intimate links between core RL theory and neurophysiology ^29–32^.

Here, we developed deep RL (DRL) models trained end-to-end, and used simulated experimental tasks from the neuroscience literature to reconcile different views on how the brain represents elapsed time. We designed and simulated novel modifications to traditional working memory tasks to investigate the extent to which task demands and task structures can affect neural representations in an ANN trained to maximize reward. We characterized the incidence and nature of these representations using neuroscience-based analyses to provide direct comparison between *in vivo* results and our *in silico* observations. We found that time-cell-esque sequences naturally emerged in recurrent DRL networks trained on working memory tasks, but so did ramping-based temporal representations. Using decoder analysis, we showed that both the time-cell-esque sequential activity and ramping activity patterns were informative about stimulus and time. Furthermore, through simulated non-mnemonic and varying-delay versions of the original task, we explored the extent to which memory and temporal dimensions of the task affected these representations. We found that memory demands are necessary for the time cells to encode information about the stimuli. In contrast, the temporal structure of the task affects both time cell encoding minimally. Thus, our models predict that time cell sequences should be present in many different tasks, even non-mnemonic tasks or tasks with different temporal structures, but the informativeness of time cell activity should depend on the memory requirements of the task. Our results provided concrete predictions for future neuroscientific investigations on how hippocampal sequences can be incorporated into cognitive map theory in the temporal dimension.

## Results

### Deep reinforcement learning agents with recurrent connections can solve a delayed nonmatch-to-sample working memory task

We simulated the mnemonic Trial-Unique, Nonmatch-to-Location (Mem TUNL) Task (***Figure 1A, Movie S1***), an episodic working memory task for which the performance in rodents has been shown to be dependent on hippocampus and the length of the delay period ^33^. We embedded the TUNL task in a 2D simulation environment resembling the touchscreen chamber commonly used in animal experiments. Agents could move up, down, left, or right, and could also “nose-poke” to interact with elements of the simulated chamber. A complete episode of the task consists of five stages: 1) At the beginning of each episode, the agent (***Figure 1A, blue square***) encounters an initiation signal (***Figure 1A, red square***), which it must navigate to and poke with in order to proceed to the sample phase. 2) During the sample phase, the sample (***Figure 1A, green squares***) will be displayed in one of two possible locations: either on the left or right corner of the triangular arena, and the agent must poke the sample. 3) Upon poking the sample, all signals are turned off, and the delay period of 40 simulation time steps starts. The end of the delay period is indicated by the onset of the initiation signal again, with which the agent must poke in order to proceed to the choice phase. 5) During the choice phase, both left and right sample locations are presented. If the agent pokes the location not displayed during the sample phase (i.e. “nonmatch”), it will receive a reward and the episode ends. This task requires the agent to remember the location of the sample throughout the delay period in order to choose correctly during the choice phase after the delay. To maximize the reward, the agent must not only remember the sample, but also navigate to desired locations in the shortest path possible without taking redundant actions.

**Figure 1.**
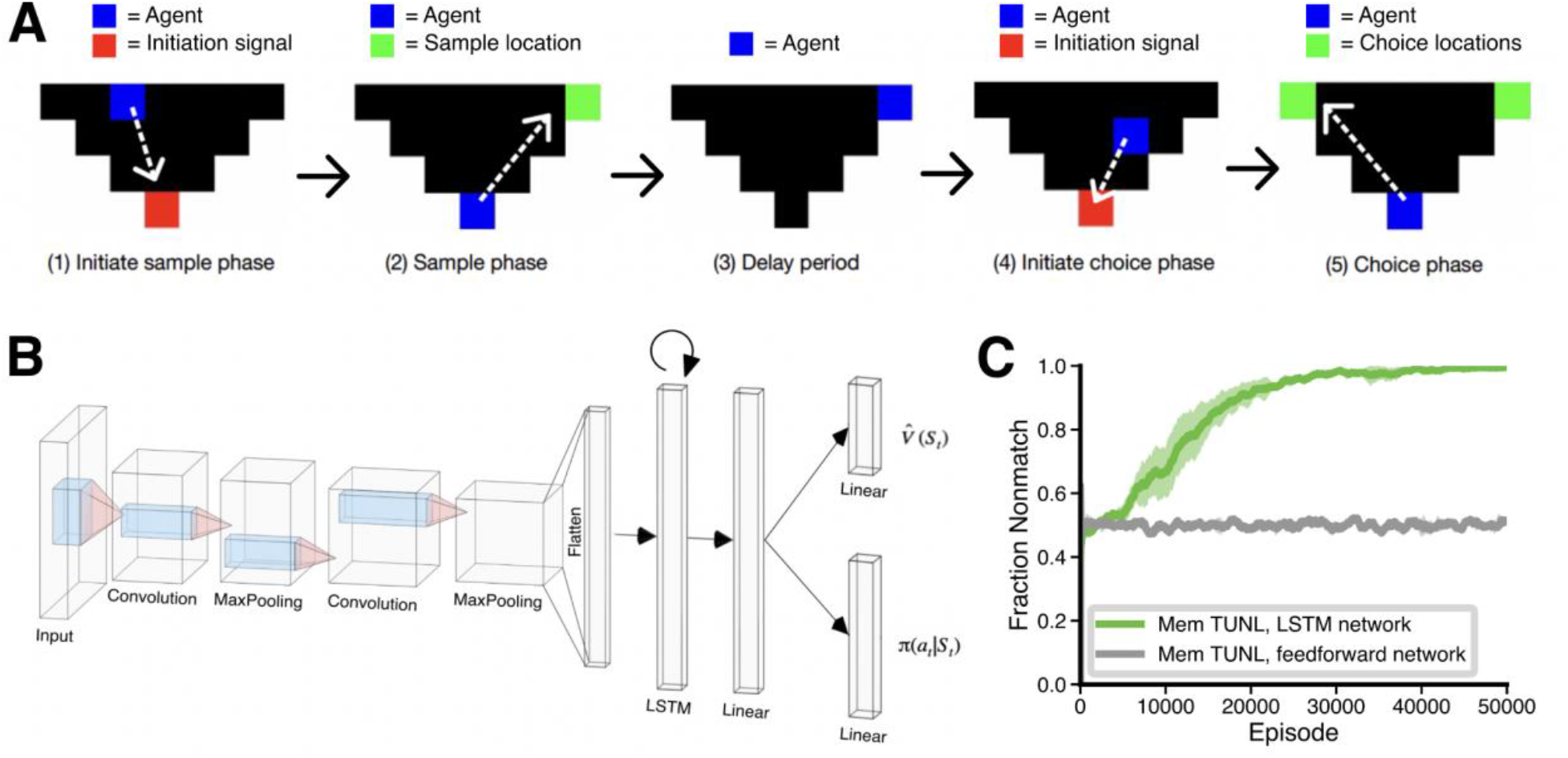
Deep reinforcement learning agent is capable of learning the simulated Trial-Unique, Nonmatch-to-Location (TUNL) working memory task. **A)** Schematic illustration of the task structure in one trial of mnemonic TUNL task (Mem TUNL). In each trial, the sample location is randomly assigned to be at either the left or right corner of the triangular arena. In this example episode, the sample location is on the right. After a delay period of 40 simulation time steps, the agent must choose the location that does **not** correspond to the sample location. In order to proceed in the task, the agent must navigate to the signal, sample, or choice locations and interact with them. Dotted white arrows show the ideal series of actions for the agent to maximize the reward. **B)** Architecture of the deep reinforcement learning (DRL) agent. At each time step, the agent receives the visual input of the state of the environment, and outputs the estimated state value 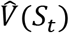 and the policy *π*(*a_t_*| *S_t_*). **C)** The agent achieves almost perfect performance on the Mem TUNL task in approximately 30000 episodes (green). In contrast, a feedforward agent chooses nonmatch at random (grey). Performance is measured as the fraction of choices that are nonmatch to sample. Solid line and shaded area represent the average and standard deviation of performance over 4 seeds, respectively.

The DRL agent used in this simulation study consisted of a visual module with a convolutional neural network, a memory module with a long short-term memory (LSTM) layer plus a linear layer, and an actor-critic RL module (***Figure 1B***; See Methods). The network was trained end-to-end with on-policy, policy gradient methods ^34^. At each simulation time step, the input provided to the agent is an RGB image of the state of the environment (i.e. a snapshot from above). The agent computes an estimated state value, and a policy according to which it will select the action in response to the current state. As noted, a LSTM was integrated into the architecture of the agent for two reasons: 1) the recurrent connectivity within LSTM cells ensures that the information about previous states can be preserved in its hidden states, providing a potential solution to our TUNL task wherein the correct choice of each trial depends on the state input received earlier in the trial; 2) the architecture of LSTM cells, which involves a gating system to direct the flow of information, avoids the vanishing and exploding gradient problem commonly observed in vanilla recurrent neural networks ^35^ and makes them an excellent algorithmic-level model of recurrent computation within the hippocampus despite the clear differences with the connectivity profile of the hippocampus ^36^.

We found that the DRL agent reliably learned to perform the mnemonic TUNL Task well (***Figure 1C, green curve***). In contrast, an agent without the recurrent LSTM component was unable to learn the task (***Figure 1C, grey curve***), verifying that the DRL agent was relying on its recurrent connections for working memory, consistent with how rodents use their hippocampus. It should be noted, though, that the DRL agent performance plateaued at a success rate that was much higher than rodents ^11,33^. This is likely because this simulated task is ultimately easier than the true task faced by the animals, and the simulated agent has no other goals or needs than solving the task. Nonetheless, the fact that the agent learns to perform this task using its recurrent connections suggests that DRL systems provide a reasonable abstract, normative model to understand the activity in biological circuits in animals performing these tasks.

### Coexistence of ramping and sequence regimes for temporal representation in LSTM cells

Currently there are two possible mechanisms by which the brain represents time that have been supported by neurophysiological recordings: the sequence regime (i.e. time cells) ^3–5,7–9^, or the ramping regime ^19–24^. Given these observations, some studies have shown the coexistence of ramping neurons and sequence neurons in the same neural circuits in the brain ^18,21,37^. However, to our knowledge, existing modelling studies of temporal representations have only focused on emulating either the ramping pattern or the sequence pattern within a population of *in silico* neurons ^38–40^. To investigate whether these regimes can emerge in task-oriented DRL agents and whether their emergence conflicts with one another, we recorded the hidden states of the 512 LSTM cells in a successfully trained DRL agent during the delay period for 1000 trials. In order to focus on cells whose activity carried temporal information, we restricted our analyses to the LSTM cells whose hidden state activities during the delay intervals across all trials had a peak-to-peak variation bigger than 10^-7^ (which selected 328/512 units, i.e. 64.06%). A ramping cell was defined as an LSTM cell whose trial-averaged temporal tuning curve monotonically increased or decreased (86/328, 26.22%; example shown in ***Figure 2B, left panel***), while the rest of the LSTM cells were defined as sequence cells (242/328, 73.78%; example shown in ***Figure 2B, right panel***), since they necessarily had a peak activation somewhere in the delay period. Importantly, these sequence cells corresponded to the time cells in the brain whose latency to their peak firing rates forms a sequence during the delay period (***Figure 2A***). Notably, both ramping cells and sequence cells carried information about the sample, since their temporal receptive fields were different under the right versus left sample conditions (***Figure 2B***).

**Figure 2.**
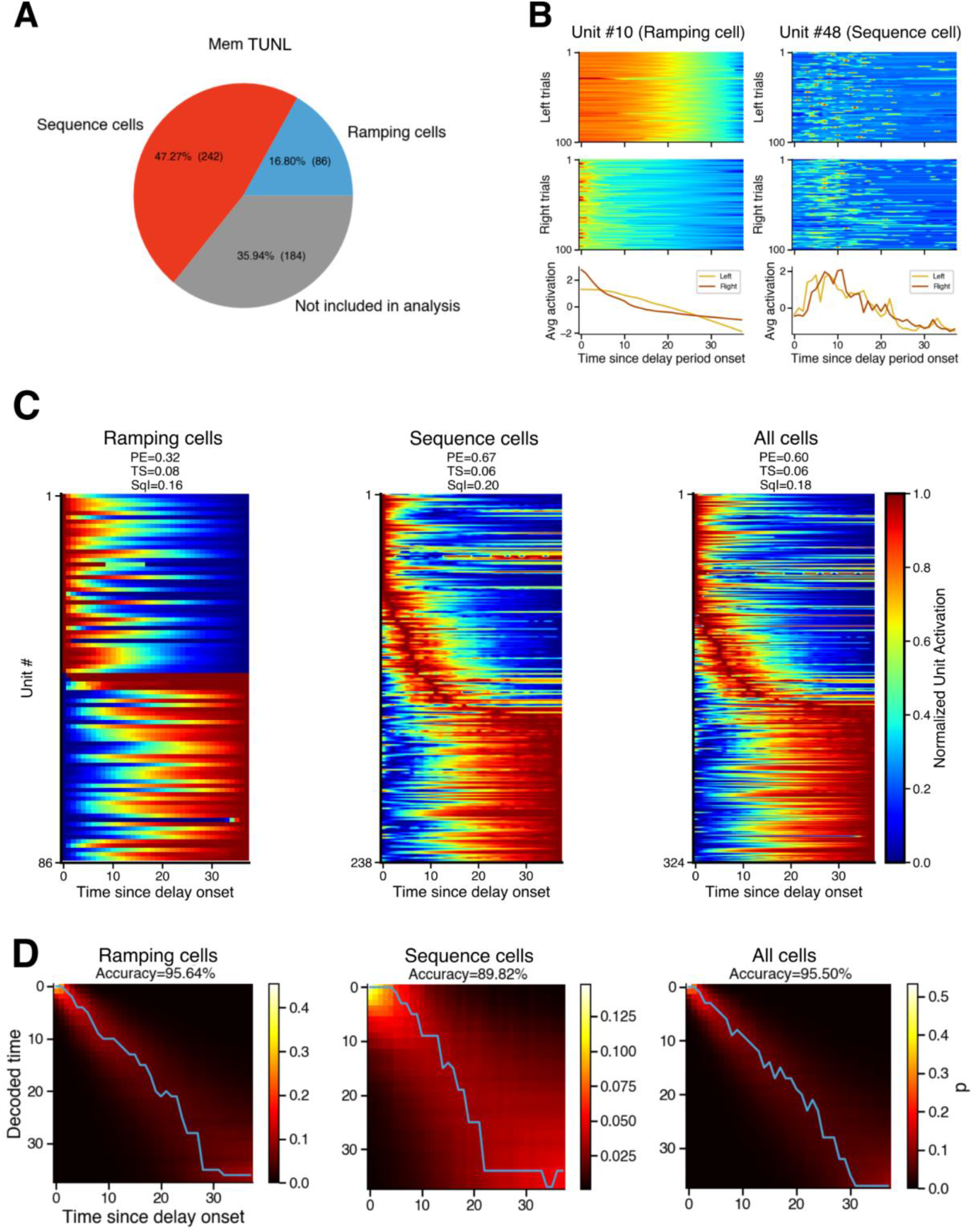
Ramping regime and sequence regime coexist in the LSTM layer during the Mem TUNL task, and both contribute to estimating the elapsed time. **A)** Pie chart showing the number and percentage of LSTM cells classified as ramping cells (blue) or sequence cells (red), or excluded from analysis due to an inadequate peak-to-peak variation in their hidden state activities (grey). **B)** An example ramping cell (left panel) and an example sequence cell (right panel). Each panel shows, from top to bottom, a heatmap of z-normalized hidden state activities during the delay period in 100 left-sample trials and 100 right-sample trials (blue and red represent the lowest and highest hidden state activities during the delay period of each trial, respectively), and the temporal tuning curves averaged over the 100 left-sample trials (yellow) and 100 right-sample trials (brown). **C)** Heatmaps showing the hidden state activities during the delay period, averaged over the 1000 recorded trials, for all ramping cells (left panel), all sequence cells (middle panel), and all analyzed cells (right panel). Each row shows the trial-averaged activities of a single cell normalized to its minimum (blue) and maximum (red). Rows in each panel are sorted by the latency to the peak hidden state activity. Peak entropy (PE), temporal sparsity (TS), and sequentiality index (SqI) for each ensemble are labeled at the top of each panel. **D)** Multi-class logistic regression decoding of elapsed time since delay onset from population hidden state activities at each time step for ramping cells (left panel), sequence cells (middle panel), and all analyzed cells (right panel). Heatmaps show probability estimates of decoded time plotted against actual elapsed time, with superimposed blue lines representing the decoded time with the highest probability estimate. Accuracy of the decoded time with the highest probability estimate is labeled at the top of each panel.

To compare the sequence cell and the ramping cell regimes at the ensemble level, we sorted the trial-averaged temporal tuning curves of cells in each regime by the latency to their peak average hidden state activity. As we expected, the tuning curves of the sequence cell population formed a continuous sequence that tiled the delay period of the TUNL task (***Figure 2C, middle panel***). Interestingly, the sequence cell ensemble was characterized by a decrease in its temporal resolution over the delay, as reflected by the overrepresentation of the beginning of the delay period as well as an increase in the width of the temporal receptive field towards the end of the delay period, which is a phenomenon commonly observed in biological time cells across brain regions and species ^3,4,9,13,17,18,41,42^. On the other hand, the average hidden state activities of different ramping cells decrease or increase at different rates (***Figure 2C, left panel***). To quantitatively characterize the dynamics observed in LSTM cells and to directly compare them to those observed in the brain, we calculated the sequentiality index (SqI) of each ensemble using the procedure described in Zhou et al.^43^ (see Methods), which takes into account 1) the entropy of the distribution of peak times (i.e. peak entropy, or PE), and 2) the sparsity of cells that peak at each given time (i.e. temporal sparsity, or TS). PE, TS, and SqI are all bounded between 0 and 1; an SqI of 1 means the dynamics of the population form a perfect sequence in which the activity of each cell in the population peaks consecutively, one-by-one at a steady pace, whereas an SqI of 0 means that either the cells did not tile time or all of the cells were active together. The results are shown at the top of each panel in ***Figure 2C***. As expected, the sequence cell regime had a higher SqI than the ramping cell regime due to a higher peak entropy. It should be noted that the SqI of LSTM sequence cells was slightly lower than that of the sequences observed in dorsolateral striatum and premotor cortex ^43^, likely due to the prolonged, sustained activity in a subset of LSTM cells towards the end of the delay period. However, whether these SqI values are different than those in the hippocampus is not yet known.

Next, we asked whether a higher sequentiality entails a better encoding of elapsed time as shown by Zhou et al.^43^. We trained a multi-class logistic regression decoder on the hidden state activities of the population of a single time step during the delay period to decode the elapsed time. We found that both ramping and sequence regimes are able to encode elapsed time in their dynamics, with slightly higher accuracies from the ramping regime (***Figure 2D***). The decoding accuracies for both ramping cells and sequence cells decreased as elapsed time progressed, reflecting the decrease in the temporal resolution observed in the population tuning curves. Altogether, our results showed that both the sequence regime and the ramping regime for temporal representation can naturally emerge in the same group of neurons in an ANN trained to perform a working memory RL task. This suggests that both time cells and ramping cells are naturally emergent phenomena in recurrent circuits of the brain.

### The emergence of stimulus-encoding sequences depends on cognitive demands

Previous research on rodent hippocampal time cells has led to disputes on whether these sequential activity patterns indeed contribute to stimulus-encoding in memory, or are merely an artifact ^8,9,26^. Here, we addressed these discrepancies and investigated the effect of task demand on temporal representations by developing a non-mnemonic version of the TUNL task (NoMem TUNL) wherein, during the choice phase, the agent may choose either the match or the nonmatch location to receive a reward, thus eliminating the demand for the agent to remember the sample location across the delay period (***Figure 3A***). The agents trained on the NoMem TUNL task had the same architectures as those trained on the Mem TUNL task (***Figure 1B***). As expected, the agents trained in the NoMem TUNL task only chose the nonmatch location half of the time (***Sup. Figure 1A***). Parallel to the Mem TUNL experiment, after training on the NoMem task, we recorded the hidden state activities from the 512 LSTM cells in the DRL agent during the delay period of 1000 trials, and selected the LSTM cells whose hidden state activities carried temporal information (264/512 units, 51.56%). We defined ramping cells and sequence cells as described previously (***Sup. Figure 1B***).

**Figure 3.**
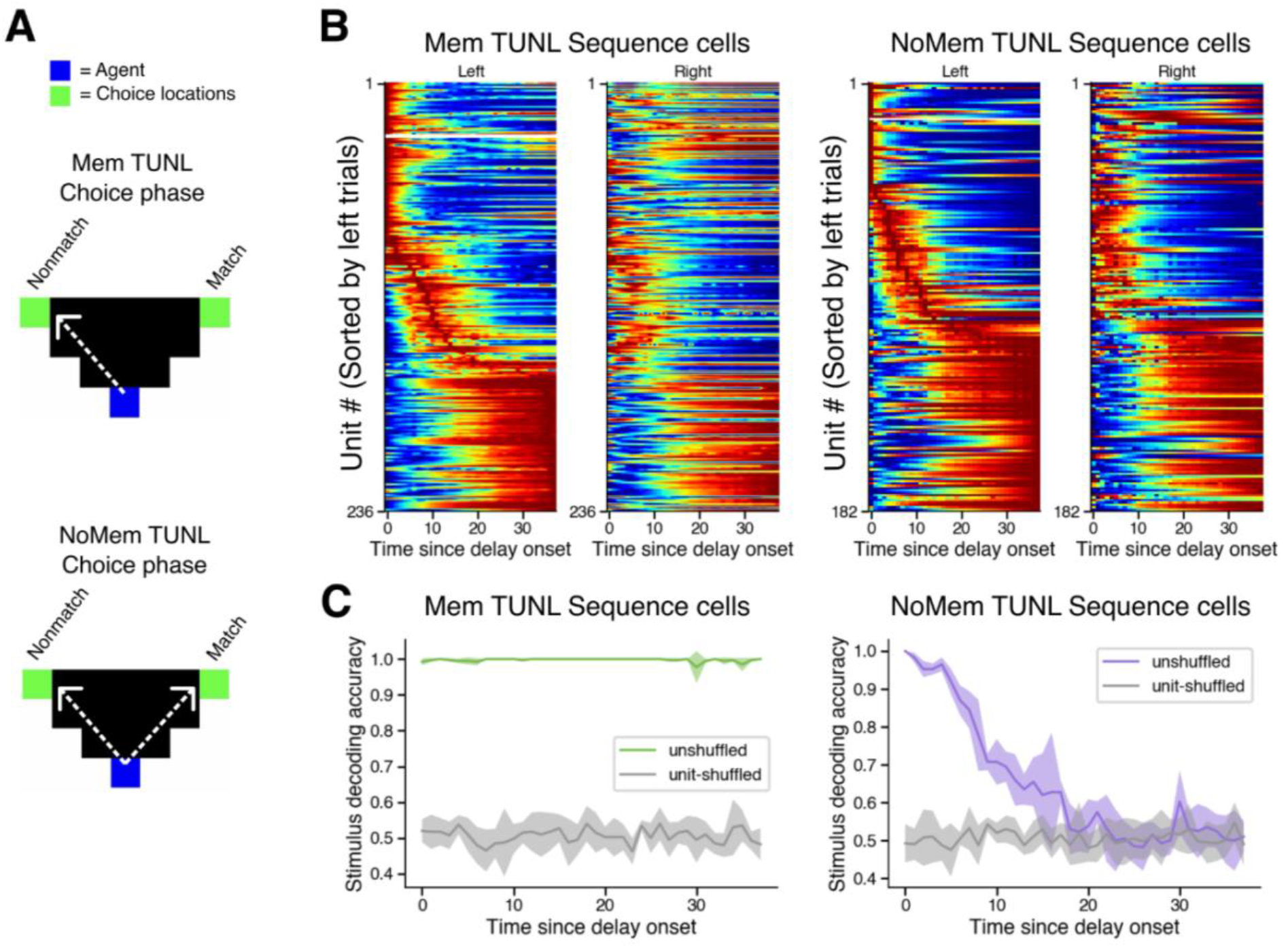
Without a demand for working memory, sequence cells progressively lose information about the sample identity over the delay period between discontiguous events. **A)** Schematic illustration of the actions (dotted white arrows) leading to a reward during the choice phase in mnemonic TUNL task (Mem TUNL, top panel) and non-mnemonic TUNL task (NoMem TUNL, bottom panel). Instead of having to select the nonmatch location, the agent may select either the nonmatch or match location in NoMem TUNL, thus eliminating the need to remember the identity of the trial-unique sample during the delay period. **B)** Sequence cell ensembles during the delay period for left-or right-sample trials under the mnemonic (left panel) or non-mnemonic (right panel) condition. Each heatmap shows the trial-averaged hidden state activities of all sequence cells, normalized to each cell’s minimum (blue) and maximum (red) activity. In all heatmaps, cells are sorted by the latency to peak hidden state activity during the left-sample trials under the corresponding task condition. **C)** SVM decoding of the sample displayed prior to the delay period from population activities at each time step during the delay period of Mem TUNL (left panel, green) and NoMem TUNL (right panel, purple). Decoding accuracy is measured by the fraction of trials decoded correctly. Decoding accuracies from unit-shuffled population activities are plotted in grey and serve as chance baseline. Solid lines and shaded areas represent mean and standard deviation of accuracies across 5 cross-validation folds.

To characterize the effect of memory demand, we first examined the temporal organization of sequence cells during different trial types under mnemonic or non-mnemonic conditions (***Figure 3B***). We found that, regardless of the presence of memory demand, left and right samples elicited different patterns of hidden state activities in the same population; in other words, the same LSTM cell would have different temporal tuning curves during the delay period after the display of different samples, as reflected in ***Figure 2B***. To quantify how informative these patterns were, we trained support vector machine (SVM) decoders to decode the identity of the sample from the hidden state activities of the sequence cell population at each single time step during the delay period on a trial-by-trial basis (we used SVMs as opposed to logistic regression here as they produced better decoding on sample states). We found that, when the agent is required to remember the identity of the sample (i.e. Mem TUNL), the sequence cell population successfully preserved the information about the sample in their hidden state activities across the entire delay (***Figure 3C, left panel, green curve***). To confirm the significance of this, we also conducted the decoding using shuffled activities (i.e. the cell identities were shuffled), which led to chance decoding performance (***Figure 3C, left panel, grey curve***). In contrast, in the absence of working memory demand (i.e. NoMem TUNL), the hidden state activities in the sequence cell population gradually lost information about the sample over the course of the delay period eventually settling at chance level (***Figure 3C, right panel***). Thus, our results lead to a specific experimental prediction: the sequential time cell regime will emerge naturally in the brain as a consequence of a delay in the task and the recurrent connections in the neural circuits regardless of the mnemonic demands; but, these time cell representations will only contribute a lasting record of sensory data from before the delay in the presence of mnemonic demands. Interestingly, we found that the ramping regime was able to decode the stimulus at any time during the delay, with or without memory demands (***Sup. Figure 1C***)

### Unscalable but informative representations of time during altered task structure

Having established that temporal representations spontaneously emerge in recurrent networks as a result of a delay period, we next asked whether changes in the temporal structure of an episode would affect the scale of temporal receptive fields and the information contained in these temporal representations. To test this, we randomly interleaved episodes with delay period durations of 20, 40, or 60 simulation time steps for both the Mem and NoMem TUNL tasks, and trained agents with the same architecture as previously described (***Figure 1B***) on these varying-delay tasks. We found that randomly changing the delay duration did not affect the learning rate of the agents (***Figure 4A***). For each task condition, after training, we again recorded the hidden state activities from the 512 LSTM cells for 1000 episodes for each delay duration, selected the LSTM cells that carried temporal information (Varying-delay Mem TUNL: 316/512 units, 61.72%; Varying-delay NoMem TUNL: 341/512 units, 66.60%), and defined ramping cells and sequence cells as described previously (***Sup. Figure 2***).

**Figure 4.**
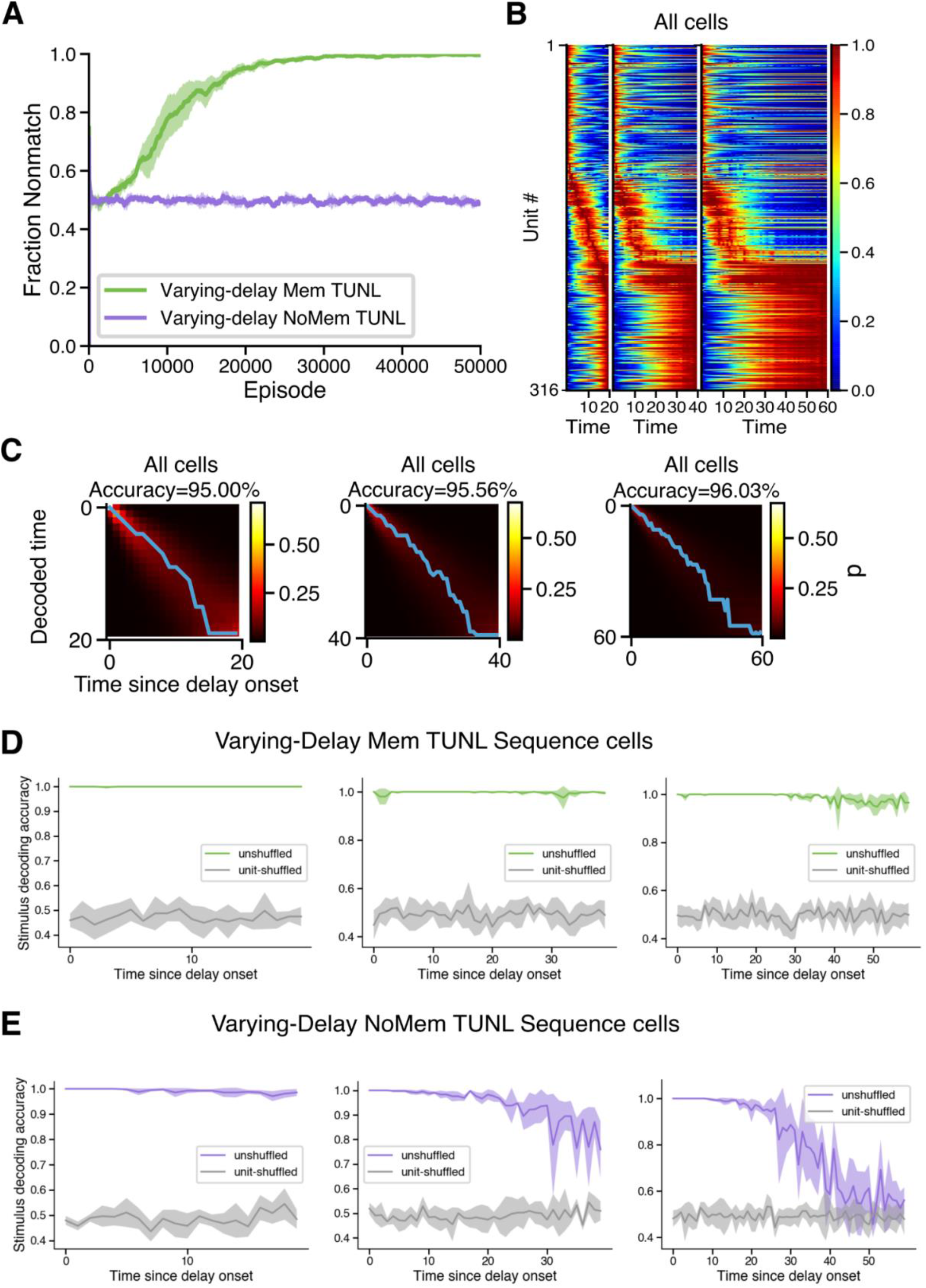
When the length of the delay abruptly changed, LSTM cells neither “retime” nor rescale, and are still able to encode time and stimulus. **A)** When the duration of the delay for each trial is randomly assigned to be either 20, 40, or 60 simulation time steps, the agent still successfully learns the Mem TUNL (green) or chooses nonmatch at random in NoMem TUNL (purple). Solid line and shaded area represent the average and standard deviation of performance over 4 seeds, respectively. **B)** LSTM cell ensembles during the delay period of 20 (left), 40 (centre), or 60 (right) time steps. Each row represents the temporal tuning curves of one LSTM cell, normalized to its minimum (blue) and maximum (red) activity in each delay duration. In all heatmaps, cells are sorted by the latency to peak hidden state activity during the 20-time step delay. **C)** Logistic regression decoding of elapsed time since delay onset from LSTM population hidden state activities at each time step for delays of length 20 (left), 40 (right) and 60 (right) time steps. Heatmaps show probability estimates of decoded time plotted against actual elapsed time, with superimposed blue lines representing the decoded time with the highest probability estimate. Accuracy of the decoded time is labeled at the top of each panel. **D)** SVM decoding of the sample from population activities at each time step during delay periods of different lengths (from left to right: 20, 40, and 60 simulation time steps) in the Mem TUNL task. Decoding accuracies from unit-shuffled population activities are plotted in grey and serve as chance baseline. Solid lines and shaded areas represent mean and standard deviation of accuracies across 5 cross-validation folds. **E)** Same as D), but for the NoMem TUNL task.

To assess how temporal representations in the LSTM cells change when the interval duration is randomly lengthened or shortened, we examined the trial-averaged temporal tuning curves of each cell in relation to the ensemble during different delay durations in Mem TUNL. We found that the temporal representations in the LSTM cells did not rescale according to the delay duration as previously reported in hippocampal time cells^44^, nor did they completely “retime” by changing their global activity patterns ^4^. Instead, LSTM cells maintained their preferences in absolute time: when the delay duration was shortened, the temporal tuning curves of LSTM cells were interrupted abruptly by the premature end of the delay; when the delay duration was lengthened, LSTM cells sustained their activities for the lengthened portion of the delay (***Figure 4B***). To investigate whether such absolute temporal representations are still informative about elapsed time, we trained a multi-class logistic regression decoder on the hidden state activities of the population of a single time step during the delay period to decode the elapsed time since delay onset. We found that the activities of LSTM cells were able to decode elapsed time with high accuracy regardless of the delay duration (***Figure 4C***). Our finding that rescaling and retiming did not occur, without impacting decodability of time, suggests that the scalability of temporal representations is not itself a prerequisite for time-encoding. It also suggests that the real brain contains additional mechanisms to support temporal calculations beyond those used in the DRL agents.

Next, to assess the effect of changes in the task structure on stimulus-encoding in the sequential “time cell” regime, we used SVM decoders to decode the identity of the sample in each trial from hidden state activities of LSTM sequence cells at each single time step during the delay period from that trial. We grouped the results based on the delay duration. We found that, in the presence of memory load (i.e. Mem TUNL), the neural activities carried information about the stimulus against the passage of time throughout the delay period regardless of the delay duration (***Figure 4D***). On the other hand, without memory demand (i.e. NoMem TUNL), stimulus-decoding accuracy of LSTM hidden state activities gradually decreased from the onset of the delay period, and continued to decrease unless interrupted by the end of the delay period (***Figure 4E***). Interestingly, comparing the stimulus-decoding accuracies for the same delay duration with or without interleaving other delay durations (***Figure 4E, middle panel; Figure 3C, right panel***), we noticed that randomly altering the delay duration slowed down the rate of information loss in the absence of memory load, possibly due to the unpredictability of the offset of the delay.

## Discussion

Inspired by the neuroscience literature on time cells and ramping cells, We trained DRL networks on simulated TUNL tasks ^33^ and examined the representations of “what happened when” in the LSTM cells ^35^. We found that the ramping regime ^19–24^ and the sequence regime^3–5,7–9^ of temporal representation emerged naturally and simultaneously in the same layer of LSTM cells (***Figure 2A-C***), and that both regimes were able to decode elapsed time and sensory stimuli (***Figure 2D, 3C, Sup. Figure 1C***). Furthermore, we simulated variations of the original TUNL task that controlled for (1) the memory demands during the delay period and (2) the temporal structure of the episodes in order to investigate the effects of task demand and task structure on the emergence and informativeness of sequence cells, which had been the subject of discrepancies in rodent studies ^8,9,26^. We found that sequence cells emerged across task demands and task structures (***Figure 3B, 4B***). However, their ability to encode sensory stimuli depended on the task demands: in the absence of memory requirements, sequence cells progressively lost information about the stimuli during the delay period (***Figure 3C, 4E***). Moreover, when we randomly altered the temporal structure of the task by lengthening or shortening the delay duration, sequence cells did not rescale or retime (***Figure 4B***), but their encoding of time and stimuli remained intact (***Figure 4C, 4D***).

Our results provide a normative framework that reconciles previous discrepancies on the consistency and informativeness of the emergence of hippocampal time cells. Consistent with Salz et al.^9^, the LSTM cells in our DRL models exhibited sequential firing activities that tiled the delay duration with or without a memory demand. We note that the non-mnemonic control task used in Salz et al. was a looping maze in which the sensory stimulus prior to the delay period did not differentiate across trials, thus was not sufficient to determine the effect of memory demand on hippocampal time cells. The fact that both Sabariego et al. ^8^ and Ahmed et al. ^26^ immobilized their rodents during the delay period forced them to conduct their analyses on cells with less stability and lower peak firing rates than previously identified hippocampal time cells in mobilized animals ^4,7,45,46^, which possibly contributed to their observations of inconsistent or uninformative sequences. The model cells used in our study differ from biological neurons in a number of ways, most notably that their activities are not measured in non-negative firing rates. To take into account this constraint, we focused on cells whose activity had large enough variations to carry temporal information, as an effort to parallel the criterion on firing rates during the selection of time cells. Nonetheless, future investigations on the interplay between hippocampal representations and memory should agree on and adopt standardized criteria for time cell identification in order to produce meaningful, comparable results.

A potential limitation of our DRL model was that, although LSTM networks have reached state-of-the-art performances on a variety of tasks that canonically require hippocampus or prefrontal cortex ^27,47,48^, it could not possibly mechanistically unify the plethora of brain regions and circuits in which time cells have been observed. For example, the temporal association memory of discontiguous events may also manifest as trace conditioning ^26,49^, for which the animal’s performance and hippocampal CA1 activities heavily rely on the input of medial entorhinal cortex layer III ^50^. On the other hand, spatiotemporal representations in hippocampus CA1 and CA3 during spatial working memory tasks ^3,7,9^ are more likely to be generated by theta sequences in medial septum ^46^. Tasks that require the animal to actively attend to the elapsed time, such as interval discrimination task or duration judgement task, often involve striatum, prefrontal and motor cortex ^14,17,21,43,51^. This may also explain lower sequentiality in LSTM cells in our models than in dorsolateral striatum and premotor cortex ^43^, as LSTM cells may not be the most suitable model for the neural dynamics of these regions.

Our results make some concrete predictions for future neurophysiological studies on hippocampal time cells. First, we predict that, without memory demands, the ability for time cells to encode sensory stimuli should gradually decrease to chance level over the delay period. Interestingly, Taxidis et al.^25^ have already observed such a progressive drop in odor-decoding accuracy in mnemonic delayed-nonmatch-to-sample task over the 7-second delay, albeit the odor-decoding accuracy remained above chance-level throughout the delay. Therefore, it would be useful for future rodent studies to measure such a “rate of forgetting” as a metric of cognitive capacity, which we predicted would benefit from randomly interleaving different delay durations. Moreover, our results predicted that, when the temporal structure of the task was altered, time cells preserved their preferences in absolute time, rather than rescaling ^44^ or retiming ^4^. The scalability of spatiotemporal representations has been theorized to contribute to the organization of cognitive maps ^52^, thus may be explained by a common circuit mechanism involving theta oscillations ^53^. Therefore, future studies on the scalability of hippocampal representations will benefit from lesion experiments to elucidate the circuitry that gives rise to scalable representations.

In summary, the results from our modelling study demonstrate that, although the emergence of temporal representations only requires the presence of a delay interval, they only meaningfully contribute to the encoding of memory under a demand for doing such. Our results link cognitive models in AI with normative models of temporal processing in neuroscience to provide concrete predictions for future experiments, and emphasize the importance of interpreting neural data under the context of the specific structure in a task.

## Methods

### Simulated TUNL task environments

The simulation environments for the mnemonic (Mem) and non-mnemonic (NoMem) TUNL tasks were designed to be compatible with the OpenAI gym framework ^54^, a suite of environments with which reinforcement learning agents interact in discrete time steps. Our environment consisted of a 4×7 inverted-triangular grid arena in which the agent freely moves, surrounded by walls of at least 1 grid thick to make up for a rectangular visual field input. The state of the environment is rendered as a coloured image, with the initiation signal in red, the left and right sample signals in green, the agent in blue, available grids in black, and the walls in white, spatially arranged according to the rodent experimental paradigm. The agent had six possible actions: move up, move down, move left, move right, interact with the signal at its current location, or stay at the same location without doing anything meaningful. When encountered with a signal, the agent not only had to move to the signal location but also interact in order to proceed in the trial. To teach the agent to move to the goal location in the shortest path possible, all actions (except for interacting with the signal when appropriate) were punished slightly.

Each trial of the Mem TUNL task consisted of five stages (***Figure 1A; Movie S1***), consistent with the rodent experimental paradigm ^33^: 1) the agent initiated the trial by interacting with the initiation signal located at the bottom of the arena; 2) during the sample phase, either the left or right signal in the top corners of the arena would be randomly displayed; 3) after interacting with the sample location, the agent experienced a signal-less delay of 40 simulation time steps, during which the agent was free to move within the arena; 4) after the delay, the agent must interact with the initiation signal again to initiate the choice phase; 5) during the choice phase, both left and right sample signals were on, and the agent must choose the one not presented during the sample phase (i.e. the nonmatch-to-sample location) to receive a reward and finish the trial. If the agent incorrectly chooses the match-to-sample location, it would receive a punishment and experience the same sample location again in the next trial as a chance to correct and learn.

The task structure of NoMem TUNL task was the same as Mem TUNL, except that, during the choice phase, the agents were allowed to select either match or nonmatch location to receive a reward and finish the trial. In the varying-delay version of the Mem and NoMem TUNL tasks, the duration of delay period for each trial was randomly selected from 20, 40, or 60 simulation time steps.

### DRL agent architectures and training details

The deep reinforcement learning (DRL) agents used in all tasks in our study (***Figure 1B***) were composed of a visual module, a memory module, and an actor-critic module. We used a deep convolutional neural network as the visual module to generate a latent representation of the visual input, a three-channel RGB image of the environment state as described in the previous section. The convolutional neural network consisted of two convolutional blocks with feature map counts of 16 and 32, respectively. Each block had a convolutional layer with kernel size 2×2 followed by max pooling with kernel size 2×2 and stride 1×1. The output of the visual module was passed to the memory module, which consisted of an LSTM layer with 512 hidden units and a linear layer with 512 hidden units. The output of the linear layer was then fed forward to a value network and a policy network ^34^, which generated an estimate of state value 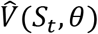 and a stochastic policy *π* (*a_t_* | *s_t_,θ*) from which the action will be sampled, respectively. We used an actor-critic algorithm, in which the network parameters were adjusted to minimize the loss *L* = *L_π_* + *L_v_*, where

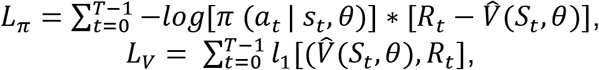

where *t* = 0,1,…,*T* - 1index the time steps in an episode with *T* experiment steps, 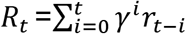 denotes the discounted return at *t*, and *l_1_* is the smooth L1 loss implemented in PyTorch ^55^.

For all tasks, we used a discount factor *γ* =0.99. The model parameters were adjusted down the gradient using Adam with *β*_1_=0.9, *β*_2_=0.999, *ϵ*=1e-8, batch size=1, and a learning rate of 1e-5.

### Neuroscience-based analysis of LSTM hidden state activities

#### Identification of ramping cells and sequence cells

We started collecting the hidden state activities from all cells in the LSTM layer after 50000 episodes of training, which was long after the performance (measured by fraction of nonmatch choices) had plateaued (***Figure 1C; Figure 4A***). For each duration of delay in each task condition, we collected the LSTM cell activities for 1000 episodes, and only kept LSTM cells whose hidden state activities during the delay interval across all recorded trials had a peak-to-peak variation larger than 10^-7^ for subsequent analysis. The remaining cells were excluded because they did not contribute significantly to meaningful representations of time, and caused numerical instability in the decoding analysis.

To determine whether the activity pattern of each LSTM cell can be characterized by the ramping regime or the sequence regime, we computed the temporal tuning curve for each cell by averaging its hidden state activity at each time during the delay across all recorded trials (***Figure 2C***). An LSTM cell was defined as a ramping cell if its trial-averaged temporal tuning curve strictly increased or strictly decreased over the duration of delay. An LSTM cell was defined as a sequence cell if it had a sufficiently large peak-to-peak variation and was not a ramping cell. We used this procedure to identify ramping cells and sequence cells in all tasks.

#### Sequentiality index

To quantitatively evaluate the extent to which the dynamics of a given ensemble of LSTM cells could be characterized by a sequence, and to provide a concrete metric for the comparison between the representations in the brain and in the DRL agents, we calculated the sequentiality index (SqI) of the sorted trial-averaged temporal tuning curves of the population as described in Zhou et al.^43^:

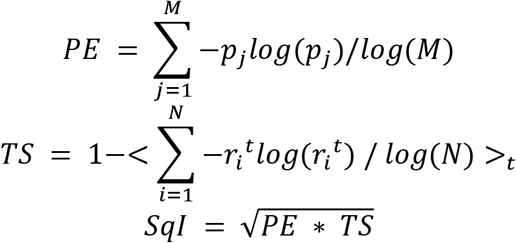

Where peak entropy (PE) measures the degree to which the peaks of the trial-averaged response of each unit in the population are evenly distributed over the delay (0<PE<1). *M*is the number of simulation time steps in the delay interval. *p_j_* is the number of cells that peak at time step *j*divided by the total number of cells in the ensemble. Temporal sparsity (TS) measures the degree to which, at each given time step, the trial-averaged responses of units are evenly distributed (0<TS<1). *N*is the number of cells in the ensemble. *r_i_^t^* is the activity of cell *i*at time *t*divided by the sum of activities of all cells at time *t*. <>_*t*_ denotes the time average. Note that, although SqI provides a characteristic metric applicable to both biological and artificial representations of time, it does not directly measure the similarity between the two.

#### Decoding of stimulus and time from single-time population activities

We used binary support vector machine (SVM) decoders to quantitatively assess how well the activities of a given population of cells at a given time during the delay period could predict the sample displayed prior to the delay (***Figure 3C; Figure 4D, 4E***). For a given population, a separate SVM decoder is constructed at each time step. We used 5-fold cross-validation to calculate the average and standard deviation of decoder accuracy. which is measured by the fraction of test trials for which the sample was decoded correctly. For each split, we held out 1 fold as testing data, and used the remaining 4 folds as training data. The hidden state activities from the given population at the given time step are z-normalized before input to the decoder to ensure that the predictions are not affected by the differences between the collective activities of a population over time or stimuli. To confirm that information about the stimulus is indeed carried by the order of cells in the population, we shuffled the order of cells in each trial at each time point, and constructed separate decoders to decode the sample from shuffled data with the same procedure above as the chance-level baseline.

We used multi-class logistic regression decoders to assess how well the collective hidden state activities of a given population at a given time point during the delay period of a given duration can decode the elapsed time since the beginning of the delay period (***Figure 2D, Figure 4C***). For each task condition and delay duration, we pooled the single-time population activities across all recorded delay periods under that task condition and delay duration, and labeled each population activity vector with the elapsed time it corresponded to. We constructed one multi-class logistic regression decoder for each pool of labeled population activity vectors, and trained it on 60% of population activity vectors selected at random. The remaining 40% of population activity vectors were used as the testing dataset: for each testing population activity vector, we calculated the probability estimates of this vector belonging to each elapsed time. We then averaged the probability estimates from testing vectors that share the same actual-time label to construct the probability heatmaps. For each actual time step, the decoded time was defined as the time giving the highest average probability estimate using population activity vectors from that actual time step as input. The accuracy of the logistic regression decoder was calculated as:

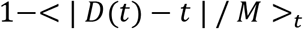

Where <>_*t*_denotes time average, *M*is the delay duration, *t*is the actual elapsed time, *D(t)* is decoded time. This ensured that calculation the accuracy of the decoded time was corrected according to the delay duration.

The simulation experiments were performed using PyTorch ^55^. All data were analyzed in Python. Decoding analyses were performed using scikit-learn packages ^56^. Additional analyses were performed using custom Python scripts, all of which are available on the author’s GitHub account^1^.

## Acknowledgements

This research is supported by grants to B.A.R from NSERC (Discovery Grant: RGPIN-2020-05105; Discovery Accelerator Supplement: RGPAS-2020-00031), Healthy Brains, Healthy Lives (New Investigator Award: 2bNISU-8), and CIFAR (Canada AI Chair; Learning in Machine and Brains Fellowship). Additionally, D.L. was supported by an IVADO Excellence Scholarship (MSc-2020-0435588519). We thank all members of the LiNC Lab for their support. In particular, we thank Chen Sun and Surya Penmetsa for helpful discussions on this project, and Annik Yalnizyan-Carson and Colleen Gillon for feedback on the manuscript.

## Author contributions

D.L. and B.A.R. conceived of the experiments. D.L. conducted the experiments and analyzed the data. B.A.R supervised the project and acquired funding. D.L. and B.A.R. wrote the manuscript.

---

**Sup. Figure 1.**
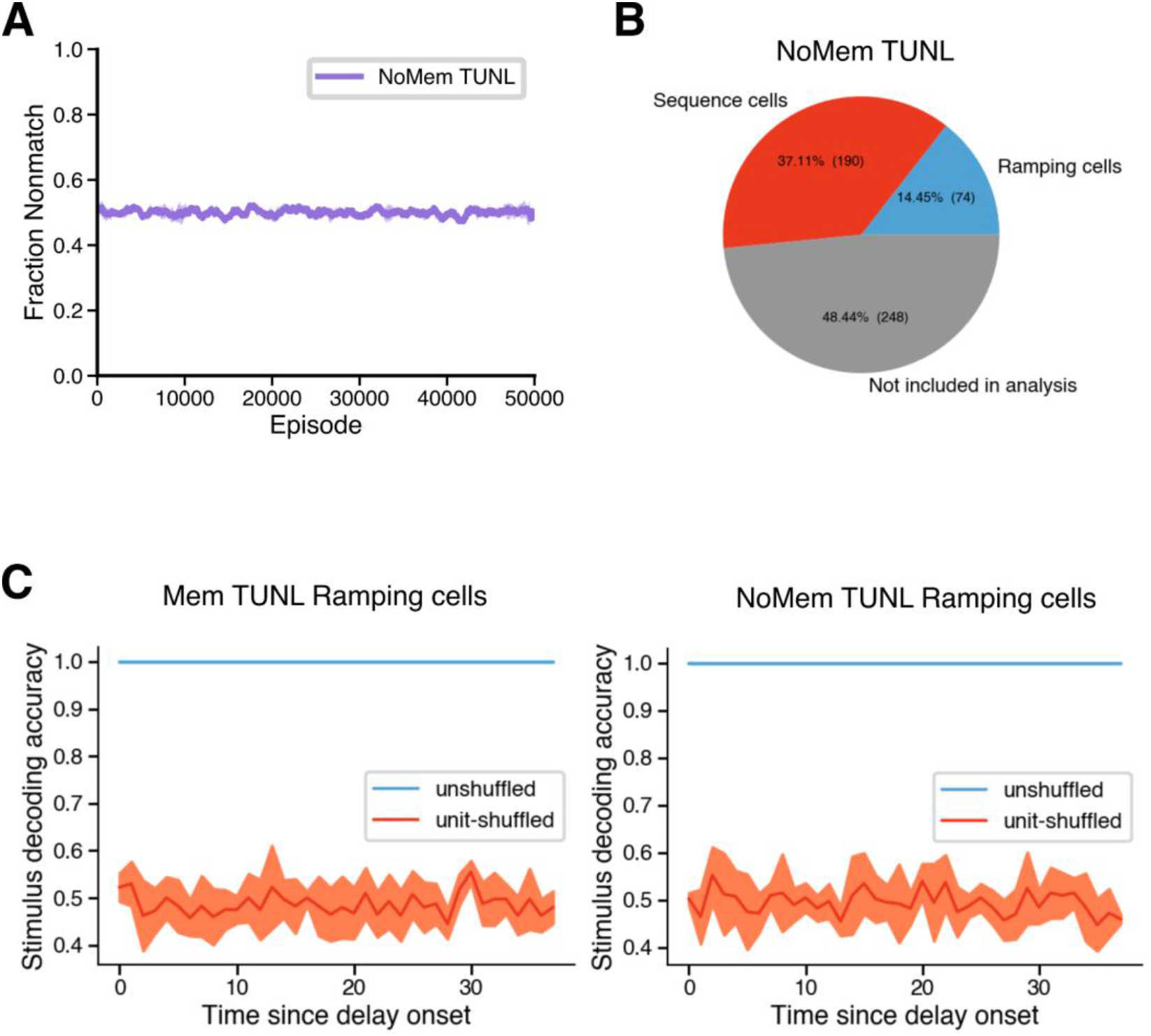
**A)** Performance of the LSTM agent on the NoMem TUNL task. **B)** Counts for sequence cells, ramping cells, or cells not included in the analysis during the NoMem TUNL task. **C)** Stimulus-decoding accuracy of the hidden state activities of ramping cells during the delay period of Mem (left) and NoMem (right) TUNL tasks.

**Sup. Figure 2.**
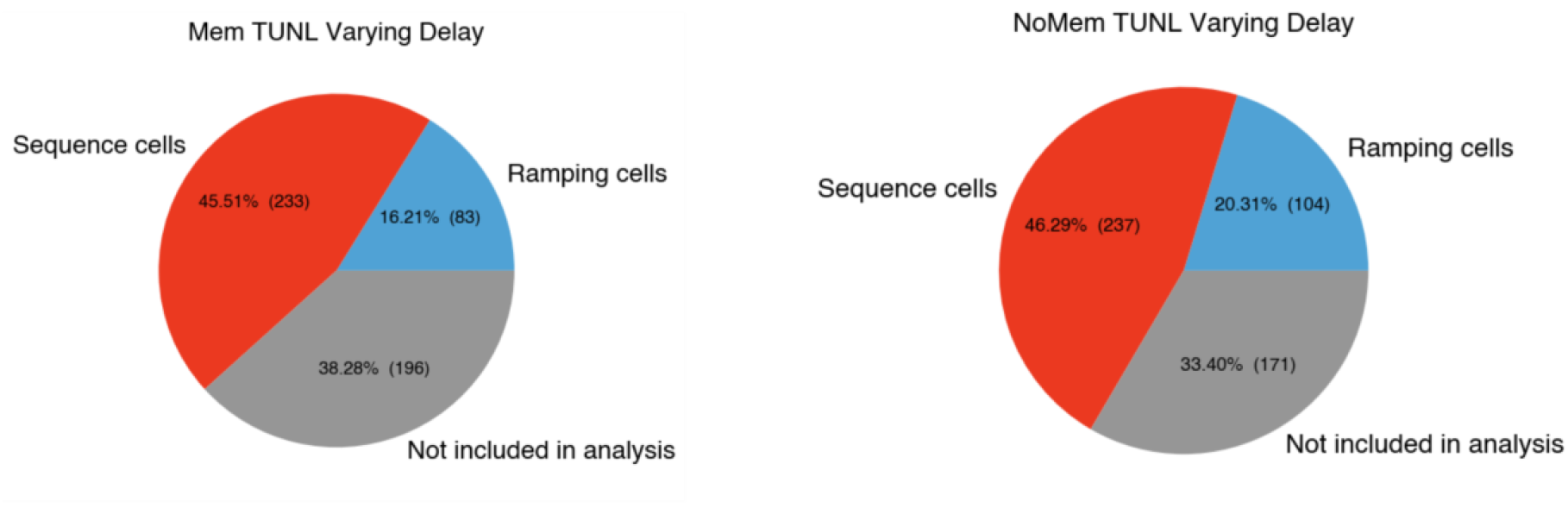
Counts for sequence cells, ramping cells, or cells not included in the analysis during varying-delay Mem (left) and NoMem (right) TUNL tasks.

## Footnotes

1 https://github.com/dongyanl1n/sim-tunl/

